# Transcriptional and Post-transcriptional Regulation of Ethylene Biosynthesis by Salicyclic Acid in Kiwifruit

**DOI:** 10.1101/2021.07.19.452972

**Authors:** Jian Wang, Xiao-fen Liu, Wen-qiu Wang, Hui-qin Zhang, Xue-ren Yin

## Abstract

Levels of ethylene, implicated in a diverse array of plants for inducing fruit ripening, is influenced by genetic and environmental factors, such as the other plant hormones. Among these, salicylic acid (SA) has been demonstrated to inhibit ethylene biosynthesis in fruit, yet the underlying regulatory mechanisms remains elusive. Here, we showed that treatment with exogenous ASA (acetylsalicylic acid) dramatically reduced ethylene production, as well as activities of ACC synthase (ACS) and ACC oxidase (ACO), in kiwifruit tissues. Comparative transcriptome analysis indicated the differential expression of ethylene biosynthetic genes (*AdACS1/2* and *AdACO5*). A screen of transcription factors indicated that AdERF105L and AdWRKY29 were ASA-responsive regulators of *AdACS1/2* and *AdACO5*, respectively. In addition to these genes, *AdACS3* and *AdACO3* were abundantly expressed in both ASA-treated and control tissues. AdACS3 protein was phosphorylated and stabilized by AdMPK16, a mitogen-activated protein kinase; while AdACO3 activity was enhanced by AdAP, an aspartic peptidase. Exogenous ASA down-regulated *AdMPK16* and *AdAP*, thereby influencing ethylene biosynthesis at a post-transcriptional level. These findings propose a multidimensional system for SA-inhibition on ethylene biosynthesis, inducing differential expression of some ethylene biosynthesis genes, as well as differential effects on protein activity on other targets.

**Summary:** Involvement of transcription factors, protein kinase and aspartic peptidase in synergistically regulating salicylic acid-induced ethylene decrease in kiwifruit flesh

## INTRODUCTION

Plant hormones are a series of small molecule compounds synthesized in plants when cells stimulated by specific signals, which are considered as important regulators of plant growth and defense (Pieterse et al., 2009; Santner et al., 2009). A long history of phytohormonal research has helped elucidate their biosynthesis, transport, signaling and response pathways, as well as the emerging underlying mechanisms of phytohormone interactions. A classic case is the cross-talk between auxin and ethylene which synergistically and antagonistically regulates different aspects of plant growth, development and fruit ripening (Stepanova et al., 2008; Yue et al., 2020; Zhang et al., 2020). Other phytohormones are also associated with ethylene to mediate plant growth and development. Exogenous gibberellin inhibits *CsACS2* and *CsACO3* to decrease ethylene production in shoot tips and regulates cucumber sexual development (Zhang et al., 2017); abscisic acid negatively modulates ethylene biosynthesis in *Arabidopsis* by transcriptionally repressing *ACS4* and *ACS8*, which is mediated by ABI4 (Dong et al., 2016); jasmonate (JA) promotes ethylene biosynthesis and fruit ripening in apple via inducing a JA signaling pathway transcription factor *MdMYC2* to activate the expression of *MdACO1* and *MdACS1* (Li et al., 2017); cytokinin and brassinosteroid enhance the stability of Type 2 ACS proteins to elevate ethylene biosynthesis in etiolated seedlings (Hansen et al., 2009). These research have builted on individual components to open new understanding of multi-phytohormone co-regulation in plants.

Salicylic acid (SA) is a key plant hormone required for inducing systemic acquired resistance (SAR) and establishing resistance to many pathogens, mainly existing in the form of free SA, SA 2-*O*-β-D-Glucoside (SAG), and methyl salicylate (MeSA) (Malamy et al., 1990; Métraux et al., 1990). The SA biosynthetic pathway has been recently clarified after the identification of three pivotal genes *SID2* (*SALICYLIC ACID INDUCTION DEFICIENT 2*), *EDS5* (*ENHANCED DISEASE SUSCEPTIBILITY 5*) and *PBS3* (*AVRPPHB SUSCEPTIBLE 3*), revealing the conversion from chorismate to SA in *Arabidopsis* (Wildermuth et al., 2001; Rekhter et al., 2019; Torrens-Spence et al., 2019). Ethylene and SA are known to synergistically accelerate plant leaf senescence and effectively coordinate plant responses to pathogens (Devadas et al., 2002; Wang et al., 2021), however there is also antagonism between them (Huang et al., 2020). Previous studies showed that ethylene signaling components ETHYLENE INSENSITIVE3 (EIN3) and ETHYLENE INSENSITIVE3-LIKE1 (EIL1) reduced SA production via negatively regulating the transcription of *SID2*, which encodes isochorismate synthase required for SA biosynthesis pathway (Chen et al., 2009). Leslie and Romani first detected decreasing ethylene production in SA-treated pear cell suspension cultures (Leslie and Romani, 1988), which was further reported in other plants, eg. Rice, kiwifruit and mungbean (Huang et al., 1993; Zhang et al., 2003; Yin et al., 2013; Khan et al., 2014). Subsequent industrial and/or research applications indicated that SA-mediated ethylene reduction is commonly associated with fruit quality, including color, softening and disease resistance (Asghari and Aghdam, 2010). However, after past three decades exposures, the underlying regulatory mechanisms of SA-inhibition of ethylene biosynthesis have not been well elucidated.

Ethylene is one of the earliest discovered phytohormones, widely involved in seed germination, leaf extension, flowering induction, organ senescence and plant response to biotic and abiotic stress (Johnson and Ecker, 1998; Bleecker and Kende, 2000). Notably, the role of ethylene as a fruit ripening inducer makes it particularly important for horticultural crops since altering its biosynthesis, perception or signaling can affect the transcription and translation of many ripening-related enzymes and proteins (Solano and Ecker, 1998; Liu et al., 2015). Ethylene in higher plants is derived from methionine (Met), and the direct precursor is 1-aminocyclopropane-1-carboxylic acid (ACC) (Adams and Yang, 1979). The ethylene biosynthetic pathway is comparitively simple: S-ade-nosylmethionine (SAM) is first converted into ACC under the catalysis of ACC synthase (ACS), and then the conversion of ACC to ethylene is achieved through ACC oxidase (ACO) (Pattyn et al., 2020). However, ethylene biosynthesis is affected by many environmental factors such as light, temperature, wounding and pathogen response (Broekaert et al., 2006; Shi et al., 2012; Zhong et al., 2012), while multiple phytohormones participate in its regulation (Hansen et al., 2009; Dong et al., 2016; Li et al., 2017; Zhang et al., 2017; Yue et al., 2020). Numerous studies have demonstrated that altering the transcription, translation and protein stability of ethylene biosynthetic enzymes, ACS and ACO, is the key step to manipulating ethylene production. For example, the WRKY family transcription factor AtWRKY33 directly binds to the promoter of *AtACS2/6* to induce expression in plant immunity (Li et al., 2012); the NAC transcription factor AtSHYG strongly transactivates the expression of *AtACO5* to regulate petiole cell expansion during root flooding (Rauf et al., 2013); mitogen-activated protein kinases MPK3/MPK6 phosphorylate two type 1 ACS isozymes (ACS2 and ACS6) to enhance their protein stability and prevent them from degradation (Liu and Zhang, 2004; Han et al., 2010; Li et al., 2012). These well-elucidated ethylene modulation pathways provide a basis to explore long-term uncharacterized regulatory links between SA and ethylene.

Kiwifruit (*Actinidia* spp.) is an ideal research plant for ethylene biosynthesis since it produces prominent ethylene and is extremely sensitive to exogenous ethylene (Zhang et al., 2018). In this study, we showed that exogenous ASA (acetylsalicylic acid) strongly inhibited ethylene production as well as enzyme activities of ACS and ACO, which are considered to be encoded by five ethylene biosynthetic genes *AdACS1/2/3* and *AdACO3/5*. Dual-luciferase assays and electrophoretic mobility shift assays (EMSA) indicated that AdERF105L and AdWRKY29 were ASA-responsive transcriptional regulators of *AdACS1/2* and *AdACO5*, respectively. In addition, we identified an aspartic peptidase protein (AdAP) which enhances protein activity of AdACO3, and verified a mitogen-activated protein kinase AdMPK16 which stabilizes AdACS3. Both of these were down-regulated by ASA treatment, resulting in enzyme activity reduction of ACS and ACO. In conclusion, these results provide a multidimensional understanding for ethylene biosynthesis and SA-ethylene antagonism.

## RESULTS

### Physiological and Biochemical Basis of SA-inhibition on Ethylene Biosynthesis

The ‘Hayward’ kiwifruit with a firmness of 62.45 N were processed into flesh discs with a diameter of 1 cm and a thickness of 2 mm. To guarantee the uniformity, each disc was cut into two equal parts for ASA treatment and control (Fig. 1A). ASA treatment dramatically inhibited ethylene production from 2.86 nL g^-1^ h^-1^ in control fruit (CK) to 0.29 nL g^-1^ h^-1^ in ASA treated fruit in the first 6 h, and similar results were detected in 12 h treated discs with a higher inhibitory efficiency (approximate 30-fold) (Fig. 1B). Exogenous ASA treatment triggered accumulation of free SA (the hydrolyzate of ASA), which increased from 0.029 ng/g to 79.80 ng/g at 6 h treatment and from 0.0599 ng/g to 183.98 ng/g at 12 h treatment, respectively (Fig. 1D).

**Figure 1.**
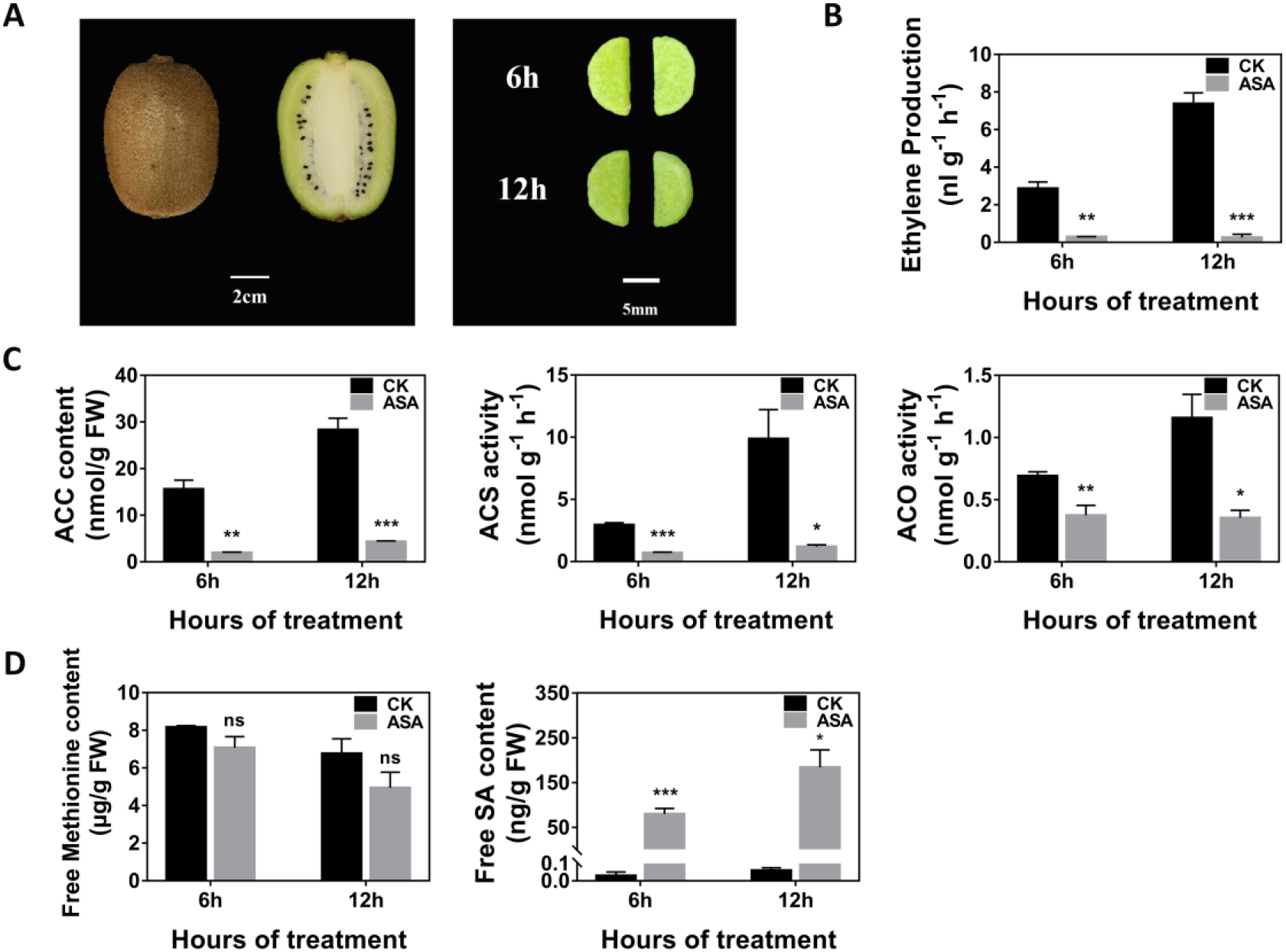
Effects of ASA treatment on kiwifruit discs. A, Fruits were processed into flesh discs with a diameter of 1 cm and a thickness of 2 mm and then cut into two equal parts for 0.5 mM acetylsalicylic acid treatments (ASA) and control (CK). Fruit discs were incubated at 28°C for 6 h and 12 h, respectively. Ethylene production (B), ethylene precursor (ACC) contents and ethylene biosynthetic enzyme (ACS and ACO) activities (C), free methionine and free SA contents (D) of kiwifruit discs were measured. Error bars indicate SEs from three replicates (*, *p* < 0.05; **, *p* < 0.01; ***, *p* < 0.001; ns, no significant).

The content of the direct ethylene precursor ACC were significantly lower in ASA treated discs (1.95 nmol/g and 4.32 nmol/g at 6 h and 12 h, respectively) than the control (15.60 nmol/g and 28.31 nmol/g at 6 h and 12 h, respectively) (Fig. 1C). In contrast, the content of methionine, the precursor of ethylene, showed no significant difference between the ASA-treated and CK discs, indicating that ethylene decrease was irrelevant to methionine content (Fig. 1D). The activities of ethylene biosynthetic enzymes (ACS and ACO) had similar trends to ACC content, which were also inhibited by ASA treatment (Fig. 1C), suggesting that ACC metabolism, involving ACS and ACO, was the main cause of ethylene reduction.

### Transcriptomic Analysis Predicted Differentially Expressed ACS/ACO Members

To investigate the key genes contributing to an ethylene decrease, RNA sequencing was applied between six mRNA libraries constructed for kiwifruit discs treated for 6 h by ASA and control, with three biological replicates. Thirty-nine Yang cycle genes and ethylene biosynthetic pathway genes were isolated by transcriptome annotation and only four gene families responded to ASA treatment (Fig. 2). The physiological and biochemical analysis (Fig. 1), guided the focuse on coding genes on ACC-related enzymes (ACS and ACO). Within the ACS family, only three members were abundant in kiwifruit discs, with *AdACS1/2* (*Achn364251/Acc05955* and *Achn339101*/*Acc30932*) being down-regulated by ASA treatment, while *AdACS3* (*Achn189421*/*Acc15646*) showed no significant difference (Fig. 2; Supplemental Fig. S1A). For the ACO family, *AdACO5* (*Achn157111*/*Acc13619*) and *AdACO3* (*Achn326461*/*Acc20538*) were more abundant than the other members, while only *AdACO5* was significantly suppressed (Fig. 2; Supplemental Fig. S1B). Recombinant proteins of AdACS1/2/3 and AdACO3/5 were purified after expression in *Escherichia coli* strain BL21 and *in vitro* protein activity assays indicated that all purified proteins had catalytic activities (Supplemental Fig. S2). Taken together, five ethylene biosynthetic genes were effective for ethylene biosynthesis in kiwifruit, while the transcriptional regulation of *AdACS1/2* and *AdACO5* could explain some of the inhibitory effect of ASA on ethylene. As the tremendous inhibitory efficiency of ASA on ethylene, the abundance of expression of *AdACS3* and *AdACO3* in both treated and control tissue suggests additional post-transcriptional regulation by ASA.

**Figure 2.**
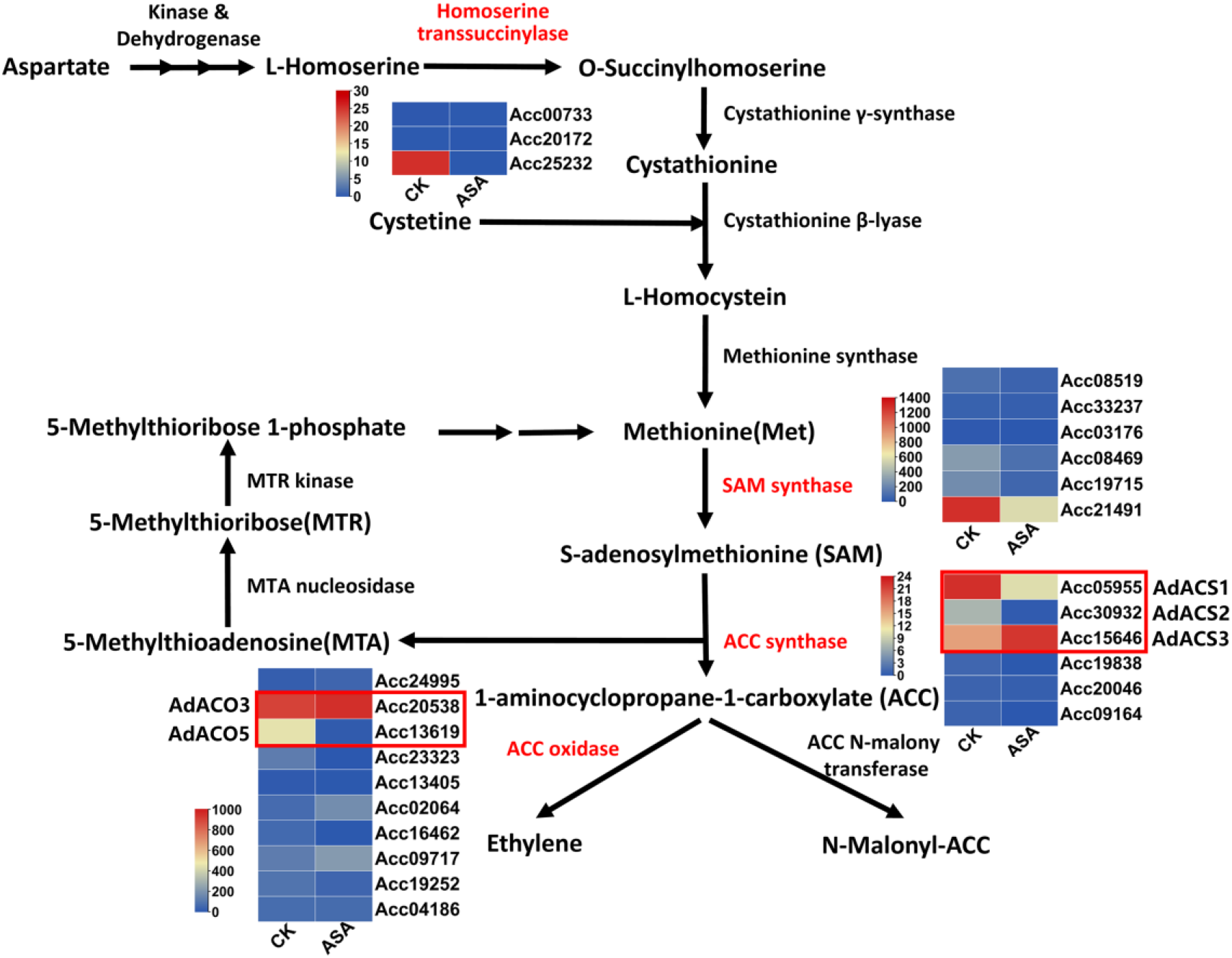
Effects of ASA treatment on expression of Yang cycle genes and ethylene biosynthetic pathway genes. Fruit discs were treated with 0.5 mM acetylsalicylic acid (ASA) or control (CK) at 28°C for 6 h and 12 h, respectively, and the comparisons were made at 6 h with three replicates at each point. The enzyme names in red are differentially expressed gene families in transcriptome data; enzyme names in black are gene families not responding to ASA. Five candidate ethylene biosynthetic genes *AdACS1/2/3* (*Achn364251*/*Acc05955*, *Achn339101*/*Acc30932* and *Achn189421*/*Acc15646*) and *AdACO3/5* (*Achn326461*/*Acc20538* and *Achn157111*/*Acc13619*) were highlighted by red boxes for further investigation.

### Transcriptional Regulation of *AdACS1/2* and *AdACO5* Promoters

The down-regulation of *AdACS1/2* and *AdACO5* could be assumed to be caused by differentially expressed transcription factors (TFs). RNA-seq results exhibited 5059 differentially expressed genes (DEGs) including 2153 up-regulated and 2906 down-regulated genes (Fold Change ≥2, FDR <0.01). These DEGs were classified into 36 groups based on two dimensions: FPKM value and absolute fold change (Fig. 3A). Two thresholds were set: 1. FPKM >200 and fold change >5 (highly expressed DEGs); 2. FPKM >30 and fold change >10 (highly differential DEGs). Based on these two selection criteria, 201 DEGs were further filtered (Fig. 3B; Supplemental Table S1). Based on gene function annotation, 201 candidate DEGs were divided into seven groups and 15 differentially expressed transcription factors (DETFs) were identified. These DETFs belong to ten family groups, including ERFs, Zinc finger proteins, and WRKYs (Fig. 3C).

**Figure 3.**
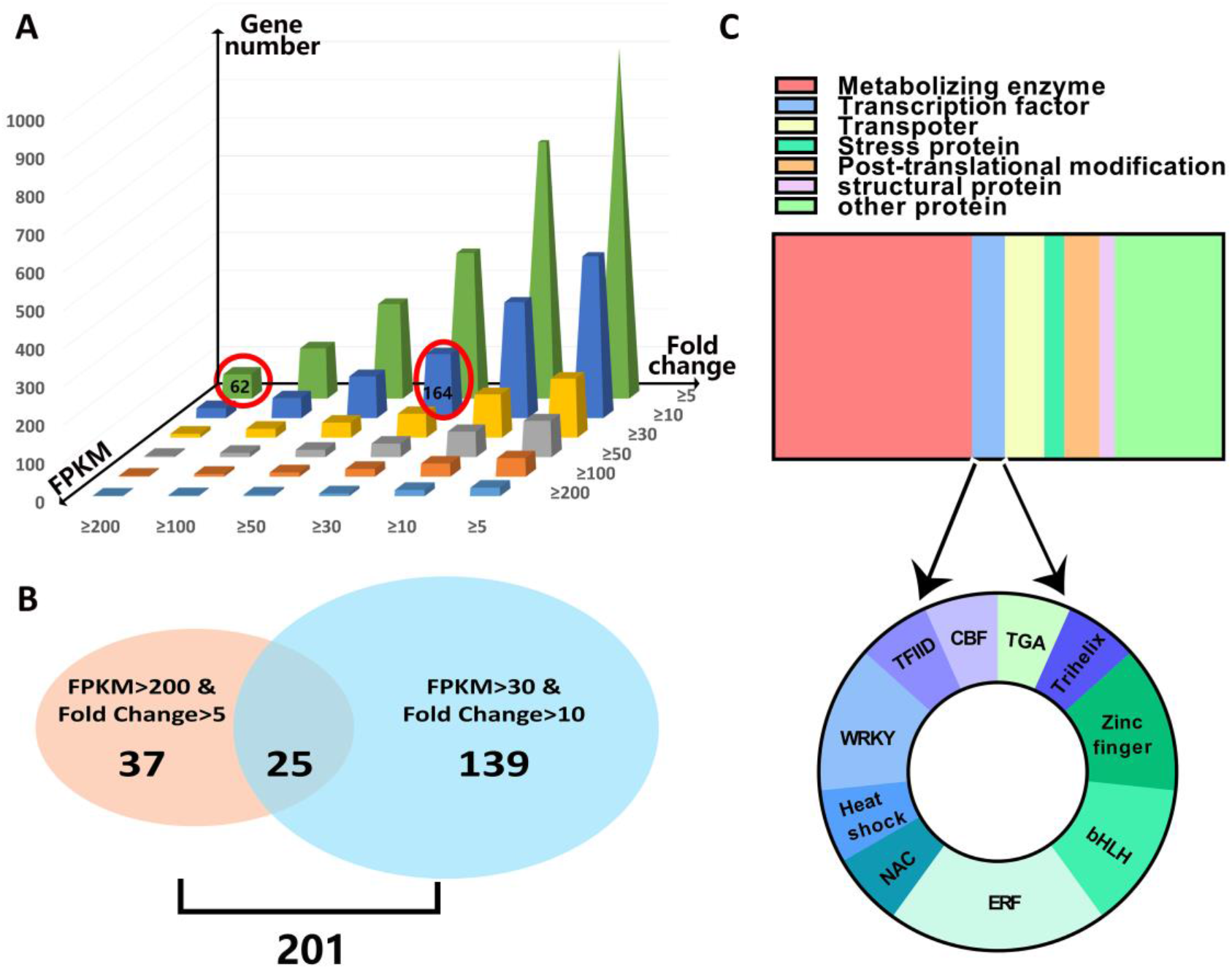
Identification of differentially expressed genes (DEGs) responding to ASA treatment by RNA-Seq analysis. Samples were analyzed with three biological replicates. A, 5059 DEGs were classified into 36 groups based on FPKM value and absolute fold change. X axis, Y axis, Z axis represent FPKM, absolute fold change, gene numbers, respectively. Two selection criteria for screening DEGs were highlighted by red circles and black numbers represent gene quantity under corresponding criteria. B, The Venn diagram showed the number of highly expressed DEGs (Pink oval) and highly differential DEGs (Blue oval), and 201 candidate genes were chosen by taking the summation of these two groups. C, 201 candidate DEGs were divided into 7 groups based on gene function annotation, and 15 differentially expressed transcription factors (DETF) were separated to verify transcriptional regulatory effects.

All of the 15 DETFs were inserted into SK vector (Hellens et al., 2005) as effectors, while promoters of *AdACS1/2* and *AdACO5* were constructed into LUC vector (Hellens et al., 2005) as reporters (Fig. 4A). Dual-luciferase assays were performed to determine the potential regulatory effects and the results showed significant transactivation by AdWRKY29 (*Achn132821*/*Acc28819*) on the *AdACO5* promoter. Meanwhile, AdERF105L (*Achn249781*/*Acc01480*) repressed the *AdACS1/2* promoter with 0.38-fold and 0.45-fold repression, respectively (Fig. 4B). RT-qPCR results indicated that *AdWRKY29* and *AdERF105L* were suppressed and induced by ASA, respectively (Fig. 4C). Besides, three *AdERF105L* over-expressed transgenic lines (Line 1/4/5) and three *AdWRKY29* over-expressed transgenic lines (Line 2/3/4) of kiwifruit plants confirmed the regulatory effects of these two TFs, via repressing or activating the expression of downstream ethylene biosynthetic genes (Supplemental Fig. S3).

**Figure 4.**
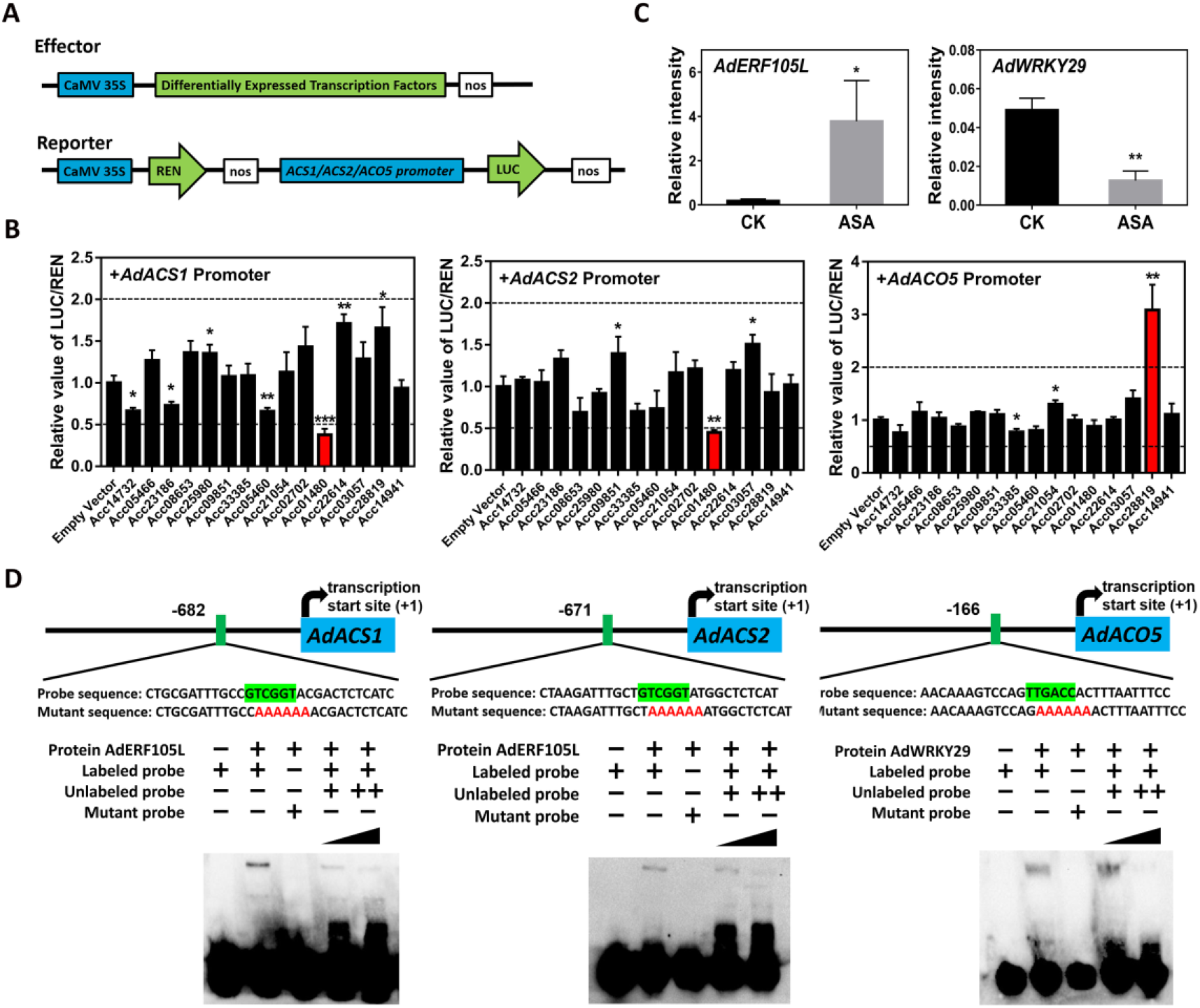
Regulatory effects of transcription factors (TF) on promoters of ASA-suppressed ethylene biosynthetic genes and analysis of binding ability. A, Schematic diagram of dual-luciferase assay. The TFs were inserted into effector vector (SK) and the promoters were constructed into reporter vector (LUC). B, Effects of 15 differentially expressed TFs on promoters of *AdACO5* and *ACS1/2*. The ratio of LUC/REN of the empty vector plus promoter was set as 1. C, Expression of two effective TFs *AdERF105L* (*Achn249781*/*Acc01480*) and *AdWRKY29* (*Achn132821*/*Acc28819*) in CK and ASA-treated kiwifruit discs. Discs were treated for 6 h. Gene expression was analyzed by RT-qPCR. Relative intensity was the expression relative to that of *AdACT*. D, Electrophoretic mobility shift assay (EMSA) showed interactions between AdWRKY29 and *AdACO5* promoter, AdERF105L and *AdACS1/2* promoter. The probe sequence used for EMSA was shown and the core binding sequences were highlighted by green box. Mutant probes were core binding sequence replaced with (AAAAAA), shown in red. Labeled probes and mutant probes were 3’ biotin-labeled; unlabeled probe was a competitor; +, ++ representing 20- and 500-fold concentration of the labeled probe, respectively. Error bars in B and C indicate SEs from five and three replicates, respectively (*, *p* < 0.05; **, *p* < 0.01; ***, *p* < 0.001).

To further investigate the physical interactions between TFs and promoters, recombinant proteins of AdWRKY29 and AdERF105L were purified and EMSA was conducted. The core binding sequence of the WRKY family is [(C/T)TGAC(C/T)] (W-box element) (Rushton et al., 2010). There were three W-box motifs in the *AdACO5* promoter (Supplemental Table S2) and only the region located between -166 bp and -161 bp showed a binding band. When mutant probe [Core binding sequence was replaced with (AAAAAA)] or two concentrations of unlabeled probe were added, the binding band disappeared or was reduced, respectively (Fig. 4D). ERF family proteins have been reported to bind the sequence of [AGCCGCC] (G-box element) and [(A/G)CCGAC] (DRE element) (Ohme-Takagi and Shinshi, 1995; Stockinger et al., 1997). One G-box and two DRE motifs were found in the promoters of both *AdACS1* and *AdACS2* (Supplemental Table S2), with a binding region found to be located -682 to -677 bp in *AdACS1* promoter (DRE) and -671 to -666 bp in *AdACS2* promoter (DRE), respectively. The effects of mutant probes and unlabeled probes verified this binding (Fig. 4D).

### ASA Influenced AdACS3 Stability via Mitogen-activated Protein Kinase

Although recombinant AdACS3 protein had high ACS activity (Supplemental Fig. S2), transcriptome data and RT-qPCR showed that expression of *AdACS3* did not respond to ASA treatment (Fig. 5A; Supplemental Fig. S1A). To detect protein abundance of AdACS3, immunoblotting analysis (IB) was conducted using an antibody that specifically recognizes AdACS3, and results showed that ACS3 protein level reduced in ASA-treated discs (Fig. 5A). To investigate whether protein reduction of AdACS3 was mediated by ubiquitin-proteasome system, the ubiquitination inhibitor MG132 (carbobenzoxy-Leu-Leu-leucinal) was applied to kiwifruit discs together with ASA. The results showed that MG132 did not significantly influence the ASA-dependent ethylene decrease (Supplemental Fig. S4).

**Figure 5.**
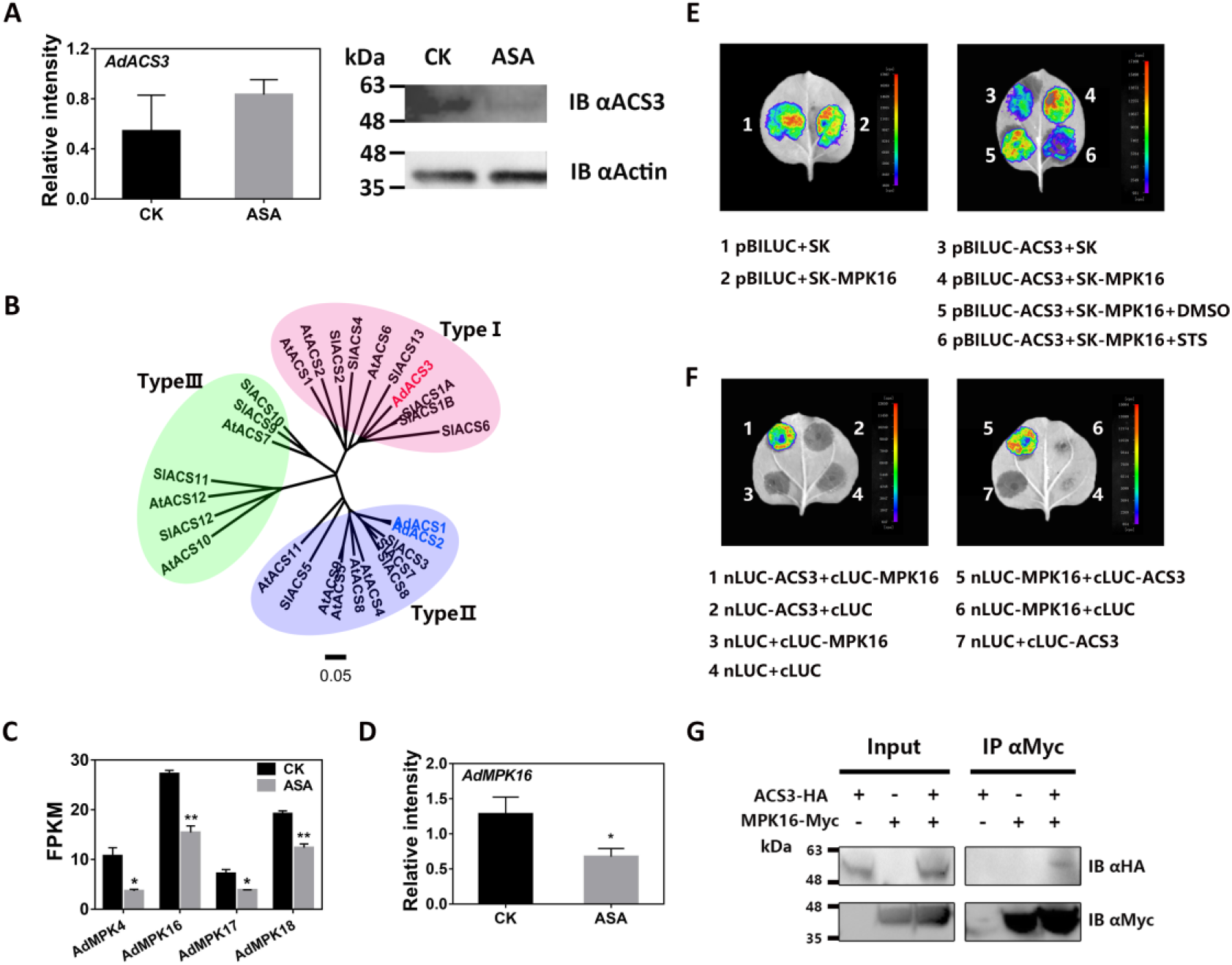
Effects of AdMPK16 on regulating phosphorylation level and protein stability of AdACS3. A, Gene exrpession and protein level of AdACS3 in CK and ASA-treated kiwifruit discs, analyzed by RT-qPCR and immunoblotting (IB), respectively. Relative intensity was the expression relative to that of *AdACT*. Protein level was confirmed by IB using ACS3 specific antibody. IB with β-actin antibody indicate similar loading. B, Phylogenetic tree analysis of ACSs in kiwifruit, *Arabidopsis* and tomato. ACSs are divided into three types based on their C terminus specificities. C, FPKM value of MPKs that were suppressed by ASA treatment. A total of 18 MPKs were found in kiwifruit genome database and four of them (*AdMPK4/16/17/18*) were down-regulated by ASA treatment. D, Gene expression of *AdMPK16* (*Achn252431*/*Acc01953*) in CK and ASA-treated kiwifruit discs, analyzed by RT-qPCR. E, Firefly luciferase imaging assays indicate that AdMPK16 stabilizes AdACS3 protein and prevents it from degradation. The strength of LUC signal (showing by different colors) indicates the protein stability of AdACS3. The empty pBILUC plus SK or AdMPK16 are negative controls showing no influence of AdMPK16 on LUC protein (1-2). STS, staurosporine, the inhibitor of protein kinase; DMSO, dimethyl sulfoxide, the solvent of STS, used as negative control. F, Interactions between AdACS3 and AdMPK16 shown by firefly luciferase complementation imaging (LCI) assays. Colored regions indicate protein-protein interactions (1, 5). Empty nLUC or cLUC plus corresponding AdAP and AdACO3 were negative controls (2-4, 6-7). G, Co-immunoprecipitation (CoIP) assay showing the interaction between AdMPK16 and AdACS3. Total protein from tobacco leaves expressing MPK16-Myc was immune-precipitated with anti-Myc antibody-conjugated agarose beads. The IP and input samples were detected by IB using HA and Myc antibodies. Error bars in A, C and D indicate SEs from three replicates, respectively (*, *p* < 0.05; **, *p* < 0.01).

Phylogenetic tree analysis indicated that AdACS3 belongs to Type 1 ACS subfamily (Fig. 5B), which contains target sites for mitogen-activated protein kinases (MAPK) at the C terminus (Pattyn et al., 2020). To explore the potential phosphorylation of AdACS3, a total of 18 MAPK genes were obtained from kiwifruit genome database and four of them (*AdMPK4/16/17/18*) were down-regulated by ASA treatment (Fig. 5C). Bimolecular fluorescence complementation (BiFC) assay showed that three MPKs (AdMPK4/16/18) could interact with AdACS3 (Supplemental Fig. S5, A and B). Firefly luciferase imaging assay showed that AdMPK16 (*Achn252431*/*Acc01953*) could stabilize the AdACS3 protein and prevent degradation, meanwhile it had no influence on empty pBILUC vector (Fig. 5E). When staurosporine (STS, the inhibitor of protein kinase) was added, the strength of LUC signal became extremely low, indicating that AdACS3 protein was phosphorylated (Fig. 5E). However, similar effects could not be observed in AdMPK4/18 infiltrated leaves (Supplemental Fig. S5C). In addition, firefly luciferase complementation imaging (LCI) and co-immunoprecipitation (CoIP) assays indicate that AdMPK16 could interact with AdACS3 protein *in vivo* and *in vitro* (Fig. 5, F and G). RT-qPCR also confirmed that *AdMPK16* was inhibited by ASA treatment (Fig. 5D). These results suggest that exogenous ASA treatment down-regulated the expression of *AdMPK16*, resulting in decreased phosphorylation levels of AdACS3 protein, and hence partially contributes to the sharp decrease in ACS enzyme activity.

### ASA Influenced AdACO3 Protein Activity via Aspartic Peptidase

Unlike AdACS3, both gene expression and protein abundance of AdACO3 was not affected by ASA treatment (Fig. 6A). To find out any potential regulators of AdACO3, a CoIP-MS assay was performed using *AdACO3* over-expressed transgenic kiwifruit plants to identify proteins with which AdACO3 forms complexes (Fig. 6B). Beads conjugated with AdACO3 specific antibody were used to immune-precipitate proteins from extracts of transgenic kiwifruit plants, with rabbit normal IgG used as a control. By removing duplicate proteins in the control, 92 unique peptides (named as ACO3 unique proteins) were found in immuno-precipitates from the extracts including AdACO3 itself by LC-MS/MS analysis (Supplemental Table S3). Combined with transcriptome data, a highly expressed and ASA-responsive gene *AdAP* (*Achn106461*/*Acc27292*) was fished (Fig. 6, C and D; Supplemental Table S3). LCI and CoIP assays confirmed the interaction between AdACO3 and AdAP (Fig. 6, E and F). In addition, the regulatory effects of AdAP on AdACO3 protein activity were detected using transient expression tobacco leaves and the results showed that AdAP did not influence gene expression or protein abundance of AdACO3 (Fig. 6G), but enhanced the enzyme activity of AdACO3 together with an increase in ethylene production compared with the control in tobacco leaves (Fig. 6H). These results indicated that AdAP increased ethylene production via enhancing the protein activity of AdACO3, while exogenous ASA treatment inhibited this process eventually leading to ethylene decrease.

**Figure 6.**
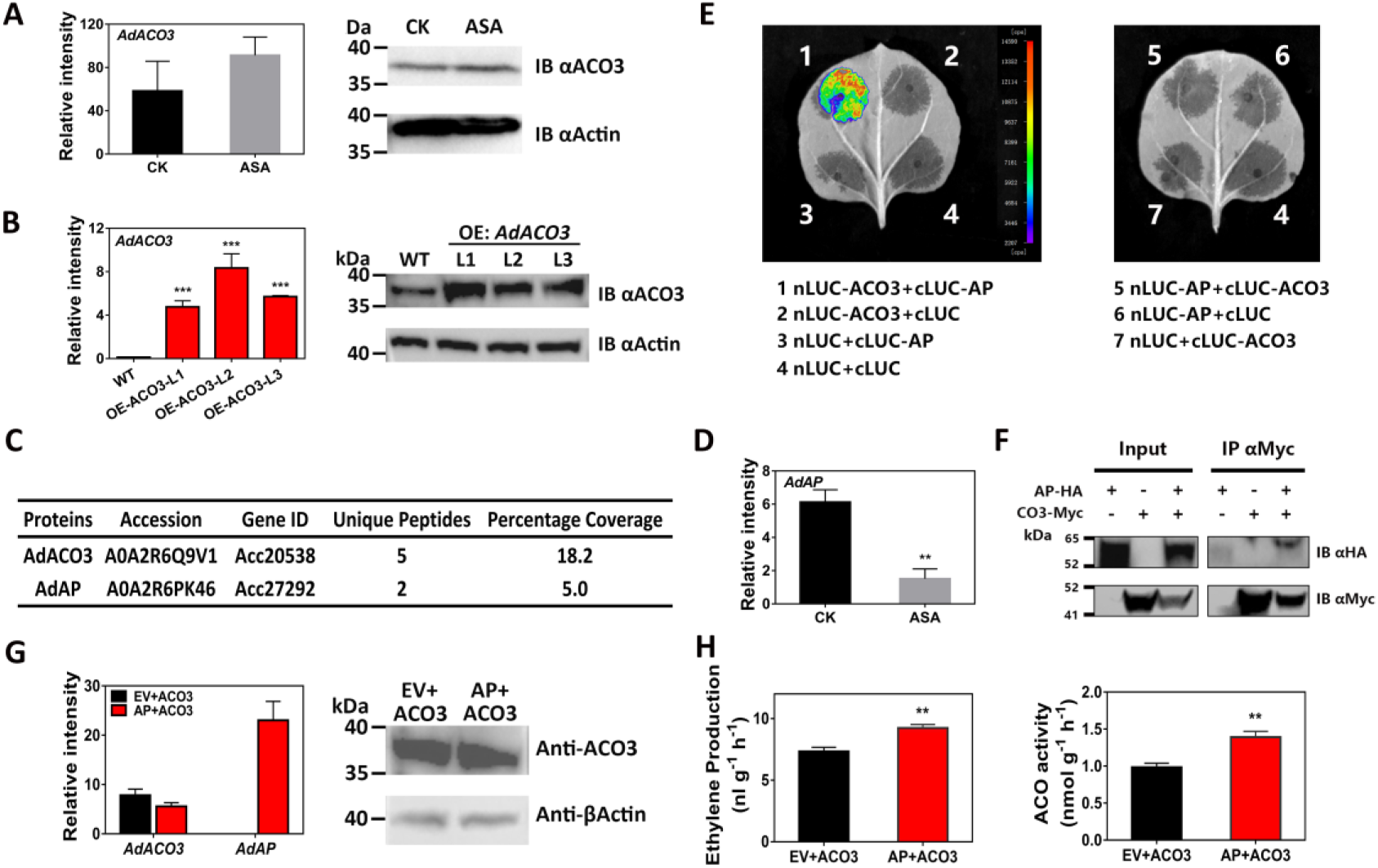
Effects of AdAP on regulating protein activity of AdACO3. Gene expression and protein abundance of AdACO3 in (A) ASA-treated kiwifruit discs and (B) *AdACO3* over-expression transgenic kiwifruit plants were shown. CK, control discs; WT, wild type. Gene expression was analyzed by RT-qPCR. Relative intensity was the expression relative to that of *AdACT*. Protein abundance was confirmed by immunoblotting (IB) using ACO3 specific antibody. IB with β-actin antibody indicates similar loading. C, AdAP protein was identified in ACO3 complexes by LC-MS/MS analysis. Beads conjugated with AdACO3 specific antibody were used to immune-precipitate (IP) proteins from extracts of transgenic kiwifruit plants and IP with rabbit normal IgG was used as a control. Accession indicates protein serial number in FASTA database. Unique Peptides represents the number of distinct peptide sequences that match the indicated protein. Percentage Coverage shows the percentage of the protein sequence covered by identified peptides. D, Gene expression of *AdAP* (*Achn106461*/*Acc27292*) in CK and ASA-treated kiwifruit discs, analyzed by RT-qPCR. E, Interactions between AdAP and AdACO3 shown by firefly luciferase complementation imaging (LCI) assays. Colored regions indicate protein-protein interactions (1). Empty nLUC or cLUC plus corresponding AdAP and AdACO3 were negative controls (2-4, 6-7). The absence of colored regions in 5 indicates that the interaction only occurred in one direction. F, Co-immunoprecipitation (CoIP) assay showing the interaction between AdAP and AdACO3. Total protein from tobacco leaves expressing ACO3-Myc was immune-precipitated with anti-Myc antibody-conjugated agarose beads. The IP and input samples were detected by IB using HA and Myc antibodies. G, Transient expressing ACO3 and AP in tobacco. ACO3 and AP were mixed a ratio of 1:1 and infiltrated into tobacco leaves. ACO3 plus empty vector was used as a control. Gene expression and protein abundance of AdACO3 in tobacco leaves were tested by RT-qPCR and IB. H, Ethylene production and ACO activity of transient infiltrated tobacco leaves. Error bars in B, D, G and H indicate SEs from three replicates (*, *p* < 0.05; **, *p* < 0.01; ***, *p* < 0.001).

### Inhibition of ASA on Ethylene Biosynthesis in Various Fruit

Four different fruit (mango, persimmon, pear and tomato) were processed into tissue discs (Fig. 7B) and treated with ASA for 6 and 12 h to test the effects on ethylene biosynthesis. All fruits used in this experiment were harvested at commercial maturity (Fig. 7A). The results showed that exogenous ASA dramatically inhibited ethylene production in these fruit discs and ethylene in ASA-treated mango and persimmon discs was undetectable at 6h (Fig. 7C). We obtained several pivotal ACS and ACO genes in these four species including *MiACS1*/*MiACO1* (Genbank no. AF170705/AJ505610), *DkACS2*/*DkACO2* (He et al., 2018), *PbACS1*/*PbACO1* (Yamane et al., 2007) and *SlACS2*/*SlACO1* (Seymour et al., 2013), and analyzed their gene expression. Except for *MiACS1*, all other genes were down-regulated by ASA treatment (Fig. 7D), which indicated a similar response to ASA on ethylene production in diverse fruit species.

**Figure 7.**
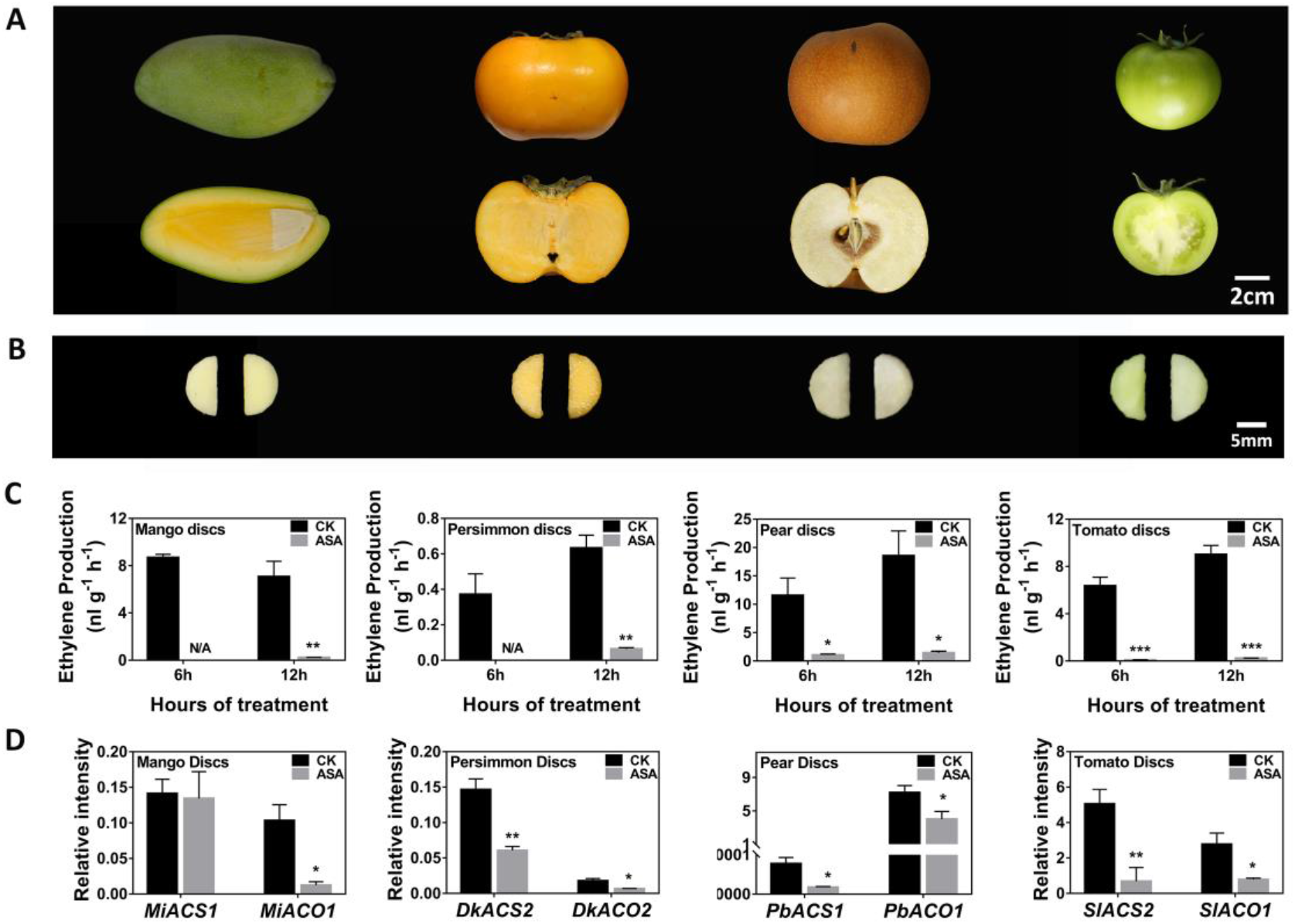
Verification of regulatory effects of ASA on ethylene biosynthesis in diverse fruit tissue discs. Four different of fruit were harvested at commercial maturity (A) and processed into discs (B). Ethylene production (C) and expression of pivotal ACSs and ACOs (D) were measured. Gene expression was analyzed by RT-qPCR. Relative intensity was the expression relative to that of *MiACT*, *DkACT*, *PbACT* and *SlACT*, respectively. Error bars in C and D indicate SEs from three replicates (*, *p* < 0.05; **, *p* < 0.01; ***, *p* < 0.001). N/A, not detected.

## DISCUSSION

Kiwifruit is a typical climacteric fruit and is extremely sensitive to exogenous ethylene. Recent studies have progressively identified biosynthetic genes, transcription factors and micro-RNAs involved in ethylene response in kiwifruit, providing a wider perspective to understand kiwifruit ripening (Zhang et al., 2018; Wang et al., 2020; Wu et al., 2020b). Multiple environmental factors and plant hormones have been confirmed to regulate kiwifruit ethylene biosynthesis, such as chilling, ozone, exogenous ethylene and MeJA (Antunes and Sfakiotakis, 2002; Minas et al., 2014; Zhang et al., 2018; Wu et al., 2020b). Here we showed that SA, a well-known phytohormone of SAR and pathogen defense, inhibited ethylene production in kiwifruit flesh discs, and similar inhibitory effects could be observed in fruit discs of other crops (Fig. 1B and 7C), as well as intact tomato fruit by injection (Supplemental Fig. S6). Despite the previous reports on the inhibitory effect of ASA and/or SA on ethylene production (Leslie and Romani, 1988; Huang et al., 1993), the investigation of the underlying mechanisms have mainly focused on the inhibition by SA on gene expression and enzyme activity (Zhang et al., 2003; Yin et al., 2013; Khan et al., 2014). Consistent with previous research, the present findings indicated that ASA inhibited ethylene production in kiwifruit can be considered to be obstructed at two steps, from SAM to ACC and from ACC to ethylene, as little change could be observed in the content of methionine (Fig. 1D). However, ACC content, as well as ACS and ACO activities, were significantly reduced (Fig. 1C).

The very unbalanced status exited for decades between the physiological, biochemical and molecular basis for ASA inhibition on ethylene production. The present results indicated the complexity of regulatory mechanisms underlying ASA affecting ethylene biosynthesis, involving both transcriptional and post-transcriptional regulation (Fig. 8). *AdACS1/2* and *AdACO5* were significantly down-regulated by ASA treatment, which was modulated by two transcription factors, AdERF105L and AdWRKY29. Among them, AdERF105L functioned as a transcriptional repressor of *AdACS1/2*, whereas AdWRKY29 was a transcriptional activator of *AdACO5.* The WRKYs belong to one of the largest plant-specific transcription factor families (Bakshi and Oelmüller, 2014), some of which are rapidly induced when plants are subjected to various stress or defense signals including SA accumulation (Eck et al., 2014). The regulatory effects of WRKYs on ethylene biosynthesis has been previously reported in wheat (Hu et al., 2018) via the repression of the promoters of *TaACS2/7/8*. Moreover, AtWRKY33 could directly activate the expression of *ACS2* and *ACS6* in response to pathogen invasion, and so play a key role in determining the kinetics and magnitude of ethylene induction in *Arabidopsis* (Li et al., 2012). The regulation of ethylene production by ERFs has also been discovered in various fruits including apple, tomato and banana (Zhang et al., 2009; Xiao et al., 2013; Li et al., 2017). Our findings indicated that ASA-inhibition of ethylene biosynthesis in kiwifruit discs was partially controlled via the transcriptional regulation of AdERF105L and AdWRKY29 on *AdACS1/2* and *AdACO5* expression (Fig. 8).

**Figure 8.**
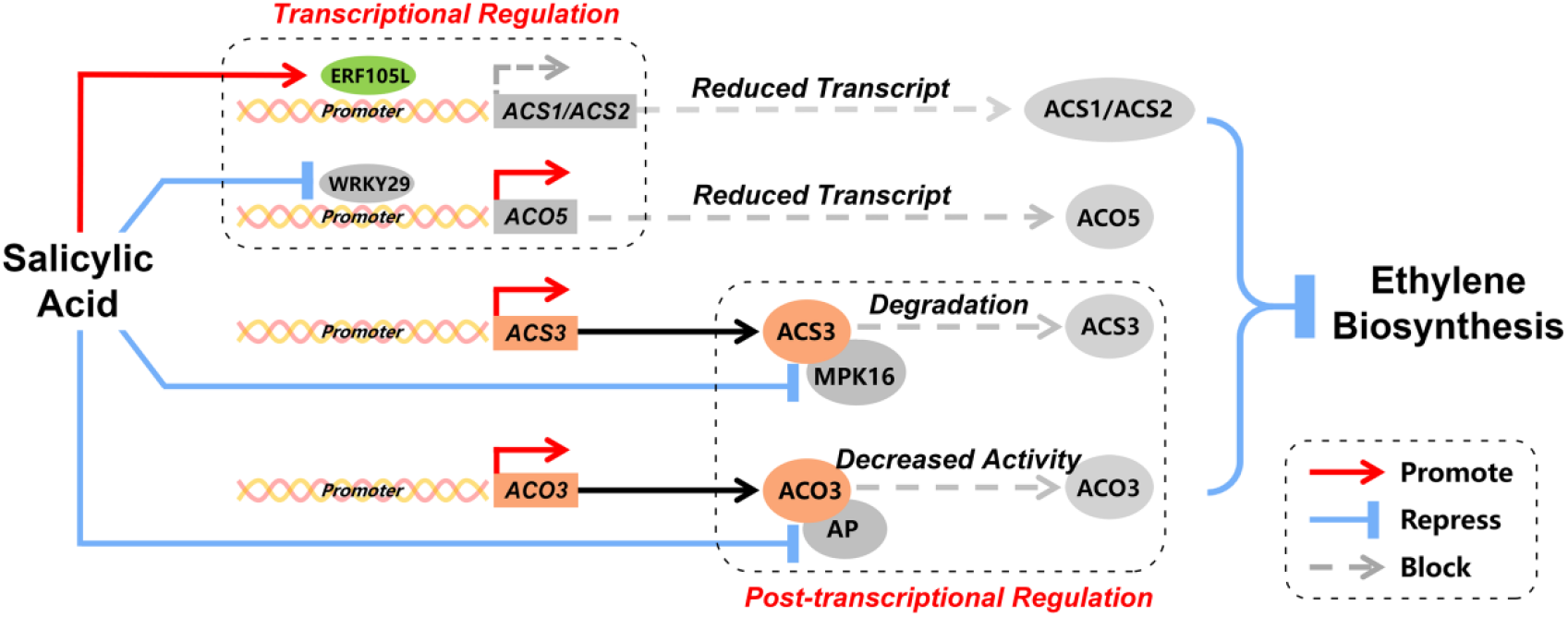
Proposed model for the mechanism by which exogenous ASA inhibits ethylene biosynthesis in kiwifruit flesh discs. AdERF105L is an ASA-induced transcriptional repressor of *AdACS1/2* and AdWRKY29 is an ASA-suppressed transcriptional activator of *AdACO5*. Both of them suppress expression of ethylene biosynthetic genes through promoter binding. Meanwhile, ASA reduces expression of *AdMPK16* and *AdAP*. *AdMPK16* encodes a protein kinase which stabilizes AdACS3 protein by increasing phosphorylation level, and *AdAP* encodes a proteolytic enzyme which enhances protein activity of AdACO3. Both of them inhibit ethylene biosynthesis at a post-transcriptional level.

As above-mentioned, the complexity of ASA regulation on ethylene production also involved post-transcriptional regulation (Fig. 8), as the expression of *AdACS3* and *AdACO3* was not altered by ASA treatment (Fig. 5A and 6A). In response to ASA treatment, AdACS3 protein levels decreased while the AdACO3 protein levels were maintained (Fig. 5A and 6A). Earlier studies have demonstrated that post-translational regulation of the ACS protein is vital for ethylene biosynthesis (Chae et al., 2003; Tan and Xue, 2014). The ACS family can be divided into three subfamilies (Pattyn et al., 2020) based on specificity of the C terminus: Type 1 with MAPK and calcium-dependent protein kinases (CDPK) target sites, Type 2 with CDPK and E3 ligases target sites and Type 3 without target sites. Phosphorylation of Type 1 ACSs has been shown to increase the ACS protein stability and thus induce ethylene production. In *Arabidopsis*, MPK3/MPK6 stabilizes ACS2/ACS6 protein *in vivo*, which greatly promotes ethylene biosynthesis when stimulated by various stresses (Liu and Zhang, 2004; Han et al., 2010). Our results indicated that the down-regulated expression of *AdMPK16* by exogenous ASA treatment reduced the phosphorylation level and promoted the degradation of AdACS3 protein, a Type 1 ACS (Fig. 5, D and E), potentially resulting in an ethylene decrease in kiwifruit discs.

The regulatory mechanism of ASA on the activity of AdACO3, which is stable and abundant at both mRNA and protein level, was revealed using the stable transgenic kiwifruit. By analyzing proteins that co-immuno-precipitated with ACO3, we identified an aspartic peptidase protein (AdAP) which exhibited high expression and significant differences between control and ASA-treated discs (Supplemental Table S3). AdAP had the function to enhance the activity of AdACO3 and to subsequently increase ethylene production (Fig. 6H). Plant aspartic proteinases (APs, also known as aspartic peptidase) are a group of proteolytic enzymes with two aspartic acid residues as catalytic active sites. The majority of typical plant APs are active in acidic pH and contain a C-terminal domain of approximate 100 amino acids named as plant specific insert (PSI) (Dunn, 2002). Typical APs have been associated with plant growth and defense including pollen germination, drought response and pathogen resistance (Xia et al., 2004; Yao et al., 2012; Huang et al., 2013). Moreover, participation of APs in plant senescence has been reported in potato and grape (Guo et al., 2013; Chen et al., 2015). However, the direct correlations between APs and ethylene production has not been previously reported. Based on our findings, it could be proposed that the formation of AP-ACO3 complexes enhances ACO3 activity to promote ethylene production. Down-regulated *AdAP* by ASA treatment reduced the complexes and inhibited ethylene biosynthesis.

## CONCLUSION

Multiple layers regulatory mechanisms of ASA inhibition of ethylene production are getting involved of both transcriptional (*AdACS1/2* and *AdACO5*) and post-transcriptional (*AdACS3 and AdACO3*) regulation, which provided the in-depth advances on this long-term existing interactions between these two plant hormones. By studying the significant inhibitory effects of ASA on ethylene production, using both simplified fruit discs, and stable transgenic plants, new regulators of the interaction between SA and ethylene have been revealed.

## MATERIALS AND METHODS

### Plant Material and Treatments

Kiwifruit (*Actinidia* spp.) were harvested from a commercial orchard in Shaanxi Province, China, in 2018, with mean total soluble solids (TSS) of 6.68%. Fruits of uniform size and free of mechanical damage were chosen and processed into flesh discs (no skin, seeds or core; 1 cm diameter, 2 mm thickness). Each disc was cut into two equal parts for reagent treatment and control experiments. The tested group was treated with 0.5 mM ASA (pH 3.5) while the control group was acidified to pH 3.5 with hydrochloric acid, both of which were supplemented with 0.4 M mannitol to maintain osmotic pressure and protect discs from rupturing or dehydration. The treatment time was 6 h or 12 h, thus 4 groups were required. Each group contained 300 half-discs, separated into three replicates of 100 half-discs each and collected in 100 mL conical flask filled with 50 mL solution. The flasks were covered with tin foil, then placed on a shaker (25 rpm) away from light and incubated at 28°C. After incubation, the discs were dried with filter paper, then 15 of them were used for ethylene measurement and the rest were frozen in liquid nitrogen and then stored at −80°C until further use.

### Ethylene Measurement

Ethylene production measurement was as described in our previous report (Yin et al., 2013; Zhang et al., 2018). The dried discs were sealed in 15 mL syringe with rubber stopper for 1 h. 1 mL of headspace gas was collected by 1 mL injector from the syringe with three replicates. The ethylene was measured with a gas chromatograph (Agilent Technologies 7890A GC System), fitted with a ProPak Q column. The temperatures of the injector, detector and oven were 140, 230 and 100°C, respectively.

### ACC Content, ACC Synthase and ACC Oxidase

The content of ACC and activities of ACS and ACO was measured following our previous reported methods (Wu et al., 2020b). Approximately 2 g of frozen discs were used for ACC extraction and 1 mL headspace gas collected from ACC reaction system was used for ACC determination. Internal standard of ACC (0.1 mM) was added to calculate the conversion efficiency of ACC to ethylene. The ACC synthase and oxidase activities were determined with 3 g and 4 g frozen discs, respectively. The ACC content and enzyme activities were analyzed by measuring ethylene production from the reaction system, using a gas chromatograph mentioned above. All the measurements were performed with three biological replicates.

### Free Amino Acids and Free SA Assays

Free amino acids were extracted from 1 g ground discs with 8 mL of 3% (w/v) sulfosalicylic acid (Shen et al., 2015) and the mixture was placed on a shaker at 200 rpm for 1 h at 4°C. The suspension was then centrifuged at 9,000 rpm for 10 min at 4°C and clarified by passage through a 0.45 μm cellulose acetate filter. Amino acids in the supernatant were further analyzed through the ninhydrin method with an automatic amino acid analyzer (L-8900; Hitachi, Tokyo, Japan).

For free SA measurement, 0.1 g frozen ground of each sample was used following the previously described protocol (Wu et al., 2007). 1 mL ethyl acetate spiked with 50 ng of D4-SA, used as the internal standard for SA, was added to each sample and high-speed oscillated on digital shaking drybath overnight at 4°C. The suspension was then centrifuged at 12,000 rpm for 10 min at 4°C and supernatants were collected in fresh 2 mL tubes. Each pellet was re-extracted with 0.5 mL of ethyl acetate with internal standard and centrifuged; supernatants were combined and evaporated to dryness on rotary evaporator at room temperature for 2 h. Then 0.5 mL of 70% chromatographic methanol was added to dissolve and mix the residue. After centrifugation at 12,000 rpm for 10 min, the supernatants were collected in liquid phase vials and then analyzed by HPLC.

### RNA Extraction and RNA-seq

Total RNA and genomic DNA was extracted by the cetyltrimethylammonium bromide (CTAB) method (Zhang et al., 2018; Wang et al., 2020). 1 μg of RNA extracted from ASA-treated and control discs with three replicates were sent for RNA-seq. Sequencing libraries were generated using NEBNext UltraTM RNA Library Prep Kit for Illumina (NEB, USA) and sequenced on an Illumina Hiseq Xten platform. Clean data was obtained by removing reads containing adapter, reads containing ploy-N and low quality reads from raw data and then mapped to the cv. Hong Yang (*Actinidia chinensis*) genome database by Tophat2 tools soft. Quantification of gene expression levels was estimated by fragments per kilobase of transcript per million fragments mapped (FPKM). The resulting *P* values were adjusted for controlling the false discovery rate and genes with an adjusted *P*-value < 0.05 found by DESeq were assigned as differentially expressed.

### cDNA Synthesis and RT-qPCR

For cDNA synthesis, gDNA removal and first-strand cDNA synthesis were conducted by PrimeScriptTM RT reagent Kit (Takara, Beijing, China) with 0.5 μg total RNA for each sample. For RT-qPCR, LightCycler^®^ 480 (Roche) and LightCycler^®^ 480 SYBR Green I Master (Roche) were used. The reaction systems and procedures were performed as previously described (Wang et al., 2020). Primers for RT-qPCR were designed by Primer3 (v.0.4.0; http://bioinfo.ut.ee/primer3-0.2324.0/) and verified by melting curve analysis. Kiwifruit *actin*, mango *actin*, persimmon *actin*, pear *actin* and tomato *actin* (Genbank no. EF063572, AF246288, AB473616, GU830958, and AB199316) was used as housekeeping genes. Sequence of primers for RT-qPCR are listed in Supplemental Table S4.

### Dual-luciferase Assay

To investigate the regulatory effects of DETFs on promoters of candidate structure genes, the dual-luciferase assay was conducted. Full lengths of 15 selected DETFs were inserted into pGreen II 0029 62-SK vector (Hellens et al., 2005), while promoters of *AdACS1/2* and *AdACO5* were recombined to the pGreen II 0800-LUC vector (Hellens et al., 2005). Sequences of primers for vector construction are listed in Supplemental Table S5. All the constructs were transferred into *Agrobacterium tumefaciens* GV3101, then cultured and dissolved in infiltration buffer (150 mM acetosyringone, 10 mM MES, and 10 mM MgCl_2_, pH 5.6), finally adjusted to an OD_600_ of 0.75. The prepared cultures of DETFs and promoters were mixed a ratio of 10:1, and infiltrated into tobacco leaves (*Nicotiana benthamiana*). Three days after infiltration, firefly luciferase and *Renilla* luciferase were analyzed with the Dual-Luciferase Reporter Assay System (Promega). Dual-luciferase assays were conducted with five biological replicates.

### Recombinant Protein Purification and EMSA

The full length coding sequence of *AdWRKY29* and *AdERF105L* were recombined to pGEX-4T-1 vector (GE) then transformed into *Escherichia coli* strain BL21 (DE3). The full length coding sequence of *AdACS1/2/3* and *AdACO3/5* were inserted into pET-32a (Novagen) vector and then transferred into *Escherichia coli* strain BL21 (DE3). Recombinant proteins were induced by 1 mM IPTG at 16°C for 20 h, then purified by a GST-tag Protein Purification Kit (Beyotime, Shanghai, China) or His TALON Purification Kit (Takara, Beijing, China). Corresponding primers are listed in Supplemental Table S2.

The probes used for EMSA were 3’-biotin end-labeled by HuaGene (Shanghai, China) and EMSA was conducted using a LightShift Chemiluminescent EMSA kit (Thermo Fisher Scientific). Mutant probes and competition probe (20-fold and 500-fold unlabeled oligonucleotides) were used to verify the binding specificity. The probes mentioned above are shown in Figure 4d.

### Protein Extraction and Immunoblot

Protein extraction was conducted using a plant protein extraction reagent (Fdbio, Hangzhou, China). Protease inhibitor mixture was added at a ratio of 1:99 into the pre-cooled extraction buffer before used. Kiwifruit samples pooled from flesh fruit discs and tissue-cultured leaves were ground into power in liquid nitrogen, and 0.5 mL extraction buffer was used for every 0.1 g of sample. After incubation on ice for 30 min, the suspension was centrifuged at 12,000 rpm for 20 min at 4°C and supernatant was collected in a fresh tube. Protein concentration was measured by aBCA protein assay kit (Fdbio, Hangzhou, China).

Immunoblot analysis was conducted following a modified method (Seguel et al., 2018). Protein extracted from tobacco or kiwifruit were separated on 12% SDS-PAGE gels and transferred to PVDF membranes (Bio-Rad) using a highly efficient wet protein transfer system (eBlot L1; GenScript, Nanjing, China). The membranes were blocked for 2 h at room temperature in TBST solution (2 mM This-HCl, pH 7.5, 75 mM NaCl and 0.05% Tween 20) with 5% nonfat-dried milk. AdACS3 and AdACO3 protein were detected using the specific rabbit polyclonal antibody at 1:5,000 dilution which were purified from immunized rabbits (HuaBio, Hangzhou, China). HA-tagged or Myc-tagged fusion proteins were detected using the rabbit polyclonal HA or Myc antibody at 1:2,500 dilution (HuaBio; 0906, R1208). Mouse monoclonal β-Actin antibody (CWBio; CW0096M) was used at 1:2,000 dilution. The membranes were incubated with diluted primary antibody at room temperature for 2 h or at 4°C overnight. After washing in TBST (10 min, 3 times), the membranes were incubated with Goat anti-Rabbit IgG-HRP (HuaBio, HA1001) or Goat anti-Mouse IgG-HRP (HuaBio, HA1003) antibody (1:10,000, 2 h, room temperature). After washing in TBST (10 min, 3 times), immunoblotting signals were obtained using a chemiluminescence kit (Fdbio; FD8000).

### CoIP and CoIP-MS

Protein-protein interactions analyzed by Co-IP assay were performed using tobacco leaves expressing corresponding protein with HA or Myc tag. The full-length coding sequence of *AdACS3* and *AdAP* were inserted into 1300-3HA vector, and full-lengths of *AdACO3* and *AdMPK16* were inserted into 1300-4Myc vector. Sequences of primers for vector construction are listed in Supplemental Table S5. The constructs were then transformed into *Agrobacterium tumefaciens* GV3101 and infiltrated into tobacco leaves (*Nicotiana benthamiana*). Two days after infiltration, the samples were collected and frozen in liquid nitrogen for protein extraction. Protein extracts were immune-precipitated using Anti-Myc Affinity Gel (Yeasen; 20587ES03) with a few left as input. IP proteins together with input proteins were analyzed by immunoblotting with HA or Myc antibody.

Immunopreciptation of *ACO3* transgenic plants was performed using Protein A Agarose (Beyotime; P2051) with AdACO3 specific rabbit polyclonal antibody or rabbit normal IgG (Beyotime; A7016). Proteins were separated by SDS-PAGE and digested with trypsin at 37°C overnight. Further LC-MS/MS analysis and data processing were performed by APTBio (Shanghai, China).

### BiFC, LCI and Firefly Luciferase Imaging Assay

BiFC and LCI tests were conducted following the previously described protocol (Wu et al., 2020a). The full length coding sequence of *AdACS3* and *AdMPK4/16/17/18* were inserted into C-terminal and N-terminal fragments of yellow fluorescent protein (YFP) vectors. The full length coding sequence of *AdACS3*, *AdMPK16*, *AdACO3* and *AdAP* were constructed into both pCAMBIA1300-nLuc and pCAMBIA1300-cLuc vectors. All the constructs were transformed into *Agrobacterium tumefaciens* GV3101 and transiently infiltrated into tobacco (*Nicotiana benthamiana*) leaves. Two days after infiltration, for BiFC, the YFP fluorescence of tobacco leaves was imaged by a Zeiss LSM710NLO confocal laser-scanning microscope; for LCI, the luciferase activity were detected by NightSHADE LB 985. Primer sequences for BiFC and LCI were listed in Supplemental Table S5.

For firefly luciferase imaging assay, full-length *AdACS3* was constructed into pBI121-Luc vector while full length coding sequence of *AdMPK4/16/18* were constructed into SK vector, using the primers listed in Supplemental Table S5. The constructs were transferred into *Agrobacterium tumefaciens* GV3101, then cultured and dissolved in infiltration buffer and adjusted to an OD_600_ of 1. The prepared cultures of ACS3 and MPK4/16/18 were mixed a ratio of 1:10, and infiltrated into tobacco leaves (*Nicotiana benthamiana*). The luciferase activity was detected by NightSHADE LB 985 after 2 d infiltration.

### Statistical Analysis

Figures were drawn with GraphPad Prism7 and Adobe Photoshop CS6. Phylogenetic tree analysis was conducted with ClustalX and FigTree. The heatmap was drawn with TBtools. Data analysis was performed with Microsoft Excel.

## Accession Numbers

Transcriptome sequencing data are available at NCBI PRJNA730419.

## Supplemental Data

The following supplemental materials are available.

**Supplemental Figure S1** FPKM values of ACS and ACO gene families in kiwifruit discs.

**Supplemental Figure S2** Recombinant protein activities of five candidate ethylene biosynthetic structural genes

**Supplemental Figure S3** Expression of transcription factors and structural genes in wild type and transgenic lines.

**Supplemental Figure S4** Effetcs of ASA and ASA+MG132 on ethylene production in kiwifruit discs.

**Supplemental Figure S5** Bimolecular fluorescence complementation (BiFC) and firefly luciferase imaging assay of AdACS3 and MPKs.

**Supplemental Figure S6** Effects of ASA on intact mature-green tomato fruits.

**Supplemental Table S1** Expression and annotation of 201 DEGs.

**Supplemental Table S2** Promoter sequences of *AdACS1, AdACS2* and *AdACO5.*

**Supplemental Table S3** List of ACO3 unique proteins.

**Supplemental Table S4** Primers for RT-qPCR.

**Supplemental Table S5** Primers for vector construction.

## ACKNOWLEDGMENTS

We thank Dr. Andrew Allan for critical reading of the manuscript and valuable comments, and thank the team at Agricultural Experiment Station of Zhejiang University for plant care.

## Notes

1 This research was supported by the National Key Research and Development Program (2018YFD1000200), the National Natural Science Foundation of China (32072635), and the Key Research and Development Program of Zhejiang Province (2021C02015), Fruit New Varieties Breeding Project of Zhejiang Province (2016C02052-7) and the Fok Ying Tung Education Foundation (161028).

## LITERATURE CITED

Adams DO, Yang SF (1979) Ethylene biosynthesis: identification of 1-aminocyclopropane-1-carboxylic acid as an intermediate in the conversion of methionine to ethylene. Proc Natl Acad Sci USA 76: 170–174

Antunes MDC, Sfakiotakis EM (2002) Chilling induced ethylene biosynthesis in ‘Hayward’ kiwifruit following storage. Sci Hortic 92: 29–39

Asghari M, Aghdam M S (2010) Impact of salicylic acid on post-harvest physiology of horticultural crops. Trends Food Sci Technol 21: 502–509

Bakshi M, Oelmüller R (2014) WRKY transcription factors: jack of many trades in plants. Plant Signal Behav 9: e27700

Bleecker AB, Kende H (2000) Ethylene: a gaseous signal molecule in plants. Annu Rev Cell Dev Biol 16: 1–18

Broekaert WF, Delauré SL, De Bolle MFC, Cammue BPA (2006) The role of ethylene in host-pathogen interactions. Annu Rev Phytopathol 44: 393–416

Chae HS, Faure F, Kieber JJ (2003) The *eto1*, *eto2*, and *eto3* mutations and cytokinin treatment increase ethylene biosynthesis in Arabidopsis by increasing the stability of ACS protein. Plant Cell 15: 545–559

Chen H, Xue L, Chintamanani S, Germain H, Lin H, Cui H, Cai R, Zuo J, Tang X, Li H, Guo H, Zhou JM (2009) ETHYLENE INSENSITIVE3 and ETHYLENE INSENSITIVE3-LIKE1 repress *SALICYLIC ACID INDUCTION DEFICIENT2* expression to negatively regulate plant innate immunity in *Arabidopsis*. Plant Cell 21: 2527–2540

Chen HJ, Huang YH, Huang GJ, Huang SS, Chow TJ, Lin YH (2015) Sweet potato *SPAP1* is a typical aspartic protease and participates in ethephon-mediated leaf senescence. J Plant Physiol 180: 1–17

Devadas SK, Enyedi A, Raina R (2002) The *Arabidopsis hrl1* mutation reveals novel overlapping roles for salicylic acid, jasmonic acid and ethylene signalling in cell death and defence against pathogens. Plant J 30: 467–480

Dong Z, Yu Y, Li S, Wang J, Tang S, Huang R (2016) Abscisic acid antagonizes ethylene production through the ABI4-mediated transcriptional repression of *ACS4* and *ACS8* in *Arabidopsis*. Mol Plant 9: 126–135

Dunn BM (2002) Structure and mechanism of the pepsin-like family of aspartic peptidases. Chem Rev 102: 4431–4458

Eck LV, Davidson RM, Wu S, Zhao BY, Botha AM, Leach JE, Lapitan NLV (2014) The transcriptional network of *WRKY53* in cereals links oxidative responses to biotic and abiotic stress inputs. Funct Integr Genomics 14: 351–362

Guo R, Xu X, Carole B, Li X, Gao M, Zheng Y, Wang X (2013) Genome-wide identification, evolutionary and expression analysis of the aspartic protease gene superfamily in grape. BMC Genomics 14: 554

Han L, Li GJ, Yang KY, Mao G, Wang R, Liu Y, Zhang S (2010) Mitogen-activated protein kinase 3 and 6 regulate *Botrytis cinerea*-induced ethylene production in Arabidopsis. Plant J 64: 114–127

Hansen M, Chae HS, Kieber JJ (2009) Regulation of ACS protein stability by cytokinin and brassinosteroid. Plant J 57: 606–614

He Y, Li J, Ban Q, Han S, Rao J (2018) Role of brassinosteroids in persimmon (*Diospyros kaki* L.) fruit ripening. J Agric Food Chem 66: 2637–2644

Hellens RP, Allan AC, Friel EN, Bolitho K, Grafton K, Templeton MD, Karunairetnam S, Gleave AP, Laing WA (2005) Transient expression vectors for functional genomics, quantification of promoter activity and RNA silencing in plants. Plant Methods 1: 13

Hu Z, Wang R, Zheng M, Liu X, Meng F, Wu H, Yao Y, Xin M, Peng H, Ni Z, Sun Q (2018) TaWRKY51 promotes lateral root formation through negative regulation of ethylene biosynthesis in wheat (*Triticum aestivum* L.). Plant J 96: 372–388

Huang J, Zhao X, Cheng K, Jiang Y, Ouyang Y, Xu C, Li X, Xiao J, Zhang Q (2013) OsAP65, a rice aspartic protease, is essential for male fertility and plays a role in pollen germination and pollen tube growth. J Exp Bot 64: 3351–3360

Huang P, Dong Z, Guo P, Zhang X, Qiu Y, Li B, Wang Y, Guo H (2020) Salicylic acid suppresses apical hook formation via NPR1-mediated repression of EIN3 and EIL1 in Arabidopsis. Plant Cell 32: 612–629

Huang YF, Chen CT, Kao CH (1993) Salicylic acid inhibits the biosynthesis of ethylene in detached rice leaves. Plant Growth Regul 12: 79–82

Johnson PR, Ecker JR (1998) The ethylene gas signal transduction pathway: a molecular perspective. Annu Rev Genet 32: 227–254

Khan MIR, Asgher M, Khan NA (2014) Alleviation of salt-induced photosynthesis and growth inhibition by salicylic acid involves glycinebetaine and ethylene in mungbean (*Vigna radiata* L.). Plant Physiol Biochem 80: 67–74

Leslie CA, Romani RJ (1988) Inhibition of ethylene biosynthesis by salicylic acid. Plant Physiol 88: 833–837

Li G, Meng X, Wang R, Mao G, Han L, Liu Y, Zhang S (2012) Dual-level regulation of ACC synthase activity by MPK3/MPK6 cascade and its downstream WRKY transcription factor during ethylene induction in Arabidopsis. PLoS Genet 8: e1002767

Li T, Xu Y, Zhang L, Ji Y, Tan D, Yuan H, Wang A (2017) The jasmonate-activated transcription factor MdMYC2 regulates *ETHYLENE RESPONSE FACTOR* and ethylene biosynthetic genes to promote ethylene biosynthesis during apple fruit ripening. Plant Cell 29: 1316–1334

Liu M, Pirrello J, Chervin C, Roustan JP, Bouzayen M (2015) Ethylene control of fruit ripening: revisiting the complex network of transcriptional regulation. Plant Physiol 169: 2380–2390

Liu Y, Zhang S (2004) Phosphorylation of 1-aminocyclopropane-1-carboxylic acid synthase by MPK6, a stress-responsive mitogen-activated protein kinase, induces ethylene biosynthesis in Arabidopsis. Plant Cell 16: 3386–3399

Malamy J, Carr JP, Klessig DF, Raskin I (1990) Salicylic acid: a likely endogenous signal in the resistance response of tobacco to viral infection. Science 250: 1002–1004

Métraux JP, Signer H, Ryals J, Ward E, Wyss-Benz M, Gaudin J, Raschdorf K, Schmid E, Blum W, Inverardi B (1990) Increase in salicylic acid at the onset of systemic acquired resistance in cucumber. Science 250: 1004–1006

Minas IS, Vicente AR, Dhanapal AP, Manganaris GA, Goulas V, Vasilakakis M, Crisosto CH, Molassiotis M (2014) Ozone-induced kiwifruit ripening delay is mediated by ethylene biosynthesis inhibition and cell wall dismantling regulation. Plant Sci 229: 76–85

Ohme-Takagi M, Shinshi H (1995) Ethylene-inducible DNA binding proteins that interact with an ethylene-responsive element. Plant Cell 7: 173–182

Pattyn J, Vaughan-Hirsch J, Poel BVD (2020) The regulation of ethylene biosynthesis: a complex multilevel control circuitry. New Phytol 229: 770–782

Rauf M, Arif M, Fisahn J, Xue GP, Balazadeh S, Mueller-Roeber B (2013) NAC transcription factor SPEEDY HYPONASTIC GROWTH regulates flooding-induced leaf movement in *Arabidopsis*. Plant Cell 25: 4941–4955

Pieterse CMJ, Leon-Reyes A, Ent SV, Wees SCMV (2009) Networking by small-molecule hormones in plant immunity. Nat Chem Biol 5: 308–316

Rekhter D, Lüdke D, Ding Y, Feussner K, Zienkiewicz K, Lipka V, Wiermer M, Zhang Y, Feussner I (2019) Isochorismate-derived biosynthesis of the plant stress hormone salicylic acid. Science 365: 498–502

Rushton PJ, Somssich IE, Ringler P, Shen QJ (2010) WRKY transcription factors. Trends Plant Sci 15: 247–258

Santner A, Calderon-Villalobos LIA, Estelle M (2009) Plant hormones are versatile chemical regulators of plant growth. Nat Chem Biol 5: 301–307

Seguel AL, Jelenska J, Herrera-Vásquez A, Marr SK, Joyce MB, Gagesch KR, Shakoor N, Jiang SC, Fonseca A, Wildermuth MC, Greenberg JT, Holuigue L (2018) PROHIBITIN3 forms complexes with ISOCHORISMATE SYNTHASE1 to regulate stress-induced salicylic acid biosynthesis in Arabidopsis. Plant physiol 176: 2515–2531

Seymour GB, Chapman NH, Chew BL, Rose JKC (2013) Regulation of ripening and opportunities for control in tomato and other fruits. Plant Biotechnol J 11: 269–278

Shen S, Wang Y, Li M, Xu F, Chai L, Bao J (2015) The effect of anaerobic treatment on polyphenols, antioxidant properties, tocols and free amino acids in white, red, and black germinated rice (*Oryza sativa* L.). J Func Food 19: 641–648

Shi Y, Tian S, Hou L, Huang X, Zhang X, Guo H, Yang S (2012) Ethylene signaling negatively regulates freezing tolerance by repressing expression of *CBF* and type-A *ARR* genes in *Arabidopsis*. Plant Cell 24: 2578–2595

Solano R, Ecker JR (1998) Ethylene gas: perception, signaling and response. Curr Opin Plant Biol 1: 393–398

Stepanova AN, Robertson-Hoyt J, Yun J, Benavente LM, Xie DY, Doležal K, Schlereth A, Jürgens G, Alonso JM (2008) *TAA1*-mediated auxin biosynthesis is essential for hormone crosstalk and plant development. Cell 133: 177–191

Stockinger EJ, Gilmour SJ, Thomashow MF (1997) *Arabidopsis thaliana CBF1* encodes an AP2 domain-containing transcriptional activator that binds to the C-repeat/DRE, a cis-acting DNA regulatory element that stimulates transcription in response to low temperature and water deficit. Proc Natl Acad Sci USA 94: 1035–1040

Tan ST, Xue HW (2014) Casein kinase 1 regulates ethylene synthesis by phosphorylating and promoting the turnover of ACS5. Cell Rep 9: 1692–1702.

Torrens-Spence MP, Bobokalonova A, Carballo V, Glinkerman CM, Pluskal T, Shen A, Weng JK (2019) PBS3 and EPS1 complete salicylic acid biosynthesis from isochorismate in *Arabidopsis*. Mol Plant 12: 1577–1586

Wang C, Dai S, Zhang ZL, Lao W, Wang R, Meng X, Zhou X (2021) Ethylene and salicylic acid synergistically accelerate leaf senescence in *Arabidopsis*. J Integr Plant Biol 63: 823–833

Wang WQ, Wang J, Wu YY, Li DW, Allan AC, Yin XR (2020) Genome-wide analysis of coding and non-coding RNA reveals a conserved miR164-*NAC* regulatory pathway for fruit ripening. New Phytol 225: 1618–1634

Wildermuth MC, Dewdney J, Wu G, Ausubel FM (2001) Isochorismate synthase is required to synthesize salicylic acid for plant defence. Nature 414: 562–565

Wu J, Hettenhausen C, Meldau S, Baldwin IT (2007) Herbivory rapidly activates MAPK signaling in attacked and unattacked leaf regions but not between leaves of *Nicotiana attenuata*. Plant Cell 19: 1096–1122

Wu W, Wang MM, Gong H, Liu XF, Guo DL, Sun NJ, Huang JW, Zhu QG, Chen KS, Yin XR (2020a) High CO2/hypoxia-induced softening of persimmon fruit is modulated by DkERF8/16 and DkNAC9 complexes. J Exp Bot 71: 2690–2700

Wu YY, Liu XF, Fu BL, Zhang QY, Tong Y, Wang J, Wang WQ, Grierson D, Yin XR (2020b) Methyl jasmonate enhances ethylene synthesis in kiwifruit by inducing *NAC* genes that activate *ACS1*. J Agric Food Chem 68: 3267–3276

Xia Y, Suzuki H, Borevitz J, Blount J, Guo Z, Patel K, Dixon RA, Lamb C (2004) An extracellular aspartic protease functions in *Arabidopsis* disease resistance signaling. EMBO J 23: 980–988

Xiao XY, Chen JY, Kuang JF, Shan W, Xie H, Jiang YM, Lu WJ (2013) Banana ethylene response factors are involved in fruit ripening through their interactions with ethylene biosynthesis genes. J Exp Bot 64: 2499–2510

Yamane M, Abe D, Yasui S, Yokotani N, Kimata W, Ushijima K, Nakano R, Kubo Y, Inaba A (2007) Differential expression of ethylene biosynthetic genes in climacteric and non-climacteric Chinese pear fruit. Postharvest Biol Technol 44: 220–227

Yao X, Xiong W, Ye T, Wu Y (2012) Overexpression of the aspartic protease *ASPG1* gene confers drought avoidance in *Arabidopsis*. J Exp Bot 63: 2579–2593

Yin XR, Zhang Y, Zhang B, Yang SL, Shi YN, Ferguson IB, Chen KS (2013) Effects of acetylsalicylic acid on kiwifruit ethylene biosynthesis and signaling components. Postharvest Biol Technol 83: 27–33

Yue P, Lu Q, Liu Z, Lv T, Li X, Bu H, Liu W, Xu Y, Yuan H, Wang A (2020) Auxin-activated MdARF5 induces the expression of ethylene biosynthetic genes to initiate apple fruit ripening. New Phytol 226: 1781–1795

Zhang AD, Wang WQ, Tong Y, Li MJ, Grierson D, Ferguson I, Chen KS, Yin XR (2018) Transcriptome analysis identifies a zinc finger protein regulating starch degradation in kiwifruit. Plant Physiol 178: 850–863

Zhang T, Li W, Xie R, Xu L, Zhou Y, Li H, Yuan C, Zheng X, Xiao L, Liu K. (2020) CpARF2 and CpEIL1 interact to mediate auxin-ethylene interaction and regulate fruit ripening in papaya. Plant J 103: 1318–1337

Zhang Y, Chen K, Zhang S, Ferguson I (2003) The role of salicylic acid in postharvest ripening of kiwifruit. Postharvest Biol Technol 28: 67–74.

Zhang Y, Zhao G, Li Y, Mo N, Zhang J, Liang Y (2017) Transcriptomic analysis implies that GA regulates sex expression via ethylene-dependent and ethylene-independent pathways in cucumber (*Cucumis sativus* L.). Front Plant Sci 8: 10

Zhang Z, Zhang H, Quan R, Wang XC, Huang R (2009) Transcriptional regulation of the ethylene response factor LeERF2 in the expression of ethylene biosynthesis genes controls ethylene production in tomato and tobacco. Plant Physiol 150: 365–377

Zhong S, Shi H, Xue C, Wang L, Xi Y, Li J, Quail PH, Deng XW, Guo H (2012) A molecular framework of light-controlled phytohormone action in *Arabidopsis*. Curr Biol 22: 1530–1535

